# Structural and biochemical insights into HCV envelope proteins for germline-targeting vaccines

**DOI:** 10.64898/2026.04.27.721052

**Authors:** Itai Yechezkel, Hila Maymon, Ariel Tennenhouse, Fang Chen, Haneen Tarabih, Jenna Weisz, Rina Fraenkel, Sarel J. Fleishman, Mansun Law, Netanel Tzarum

## Abstract

Despite effective antivirals, Hepatitis C remains a global health burden, reflecting the difficulty of eliciting broadly neutralizing antibodies (bnAbs). For the Hepatitis C virus, bnAb responses are biased toward VH1-69–encoded Abs targeting a conserved epitope on the E2 glycoprotein, suggesting that effective immunogens must engage the corresponding germline B cell receptors. However, the structural basis of germline recognition remains unclear. Here, we show that germline-reverted VH1-69 bnAbs recognize the E2 core domain in a manner closely resembling mature Abs binding. High-resolution structures reveal a conserved mode of engagement, indicating that key epitope features are accessible prior to affinity maturation. Guided by these insights, we engineered E2 variants to enhance germline Ab interactions and stability. Although these designs increased affinity, they did not improve engagement of naïve B cells, highlighting a key constraint in germline-targeting strategies. Together, our findings define principles governing germline recognition and inform the rational design of an HCV vaccine.

## Introduction

HCV is a bloodborne pathogen mainly transmitted through the transfusion of unscreened blood products, hazardous medical procedures, needle sharing among people who inject drugs (PWIDs), and unsterilized equipment in places like tattoo parlors (Hepatitis C Surveillance | 2022 Hepatitis Surveillance | CDC). The 2025 WHO global hepatitis report estimated that 50 million individuals, roughly 0.6 % of the global population, were living with HCV in 2022. The virus accounted for approximately 1 million new infections and 242,000 deaths in 2022. While approximately 30% of acute cases resolve spontaneously within 6 months, the remaining progress to chronic hepatitis C, which frequently leads to severe complications, including liver cirrhosis in 20% of patients and at increased risk for hepatocellular carcinoma (Hepatitis C Surveillance, CDC; Wester *et al*, 2025; Hepatitis C, WHO). HCV management has been revolutionized by the introduction of direct-acting antivirals (DAAs) that target the non-structural proteins essential for HCV replication. While second-generation DAAs introduced in 2014 achieve cure rates above 95%, several systemic barriers remain, including, but not limited to, the asymptomatic nature of early HCV infection, delayed diagnosis, irreversible liver damage (Cox, 2015), and the lack of protection against reinfection (Matelski *et al*, 2025; Yeung *et al*, 2022). These factors highlight the urgent need for a prophylactic vaccine to attain global HCV eradication.

HCV is an enveloped, positive-sense, single-stranded RNA virus that belongs to the Hepacivirus genus within the Flaviviridae family (Taxon Details, ICTV; Sallam & Khalil, 2024; Yechezkel *et al*, 2021). A major challenge in developing an HCV vaccine is the virus’s extreme genetic diversity, which is currently classified into eight main genotypes and over 90 subtypes (Taxon Details, ICTV; Di Stefano *et al*, 2023). This diversity is driven by the NS5B RNA-dependent RNA polymerase, which lacks proofreading capabilities, resulting in high error rates and complex viral variant populations within a single host, enabling rapid immune evasion (Farci, 2011; Rothhaar *et al*., 2025). The envelope (Env) glycoproteins E1 and E2 form a noncovalent heterodimer on the surface of the virion and facilitate cell entry. E2 functions as the main receptor-binding protein that binds the CD81 host cell receptor along with other host factors (Kumar *et al*, 2023). Consequently, the E2 receptor-binding site serves as an accessible target for broadly neutralizing antibodies (bnAbs). bnAbs facilitate viral clearance and protection against a wide range of HCV strains, and are therefore a valuable tool for studying conserved viral targets for vaccine development. However, the E2 protein contains several variable regions and is heavily glycosylated, which shields it from host immune responses (Fig. 1A, B) (Kulakova *et al*, 2025; Lavie *et al*, 2018).

**Fig. 1:**
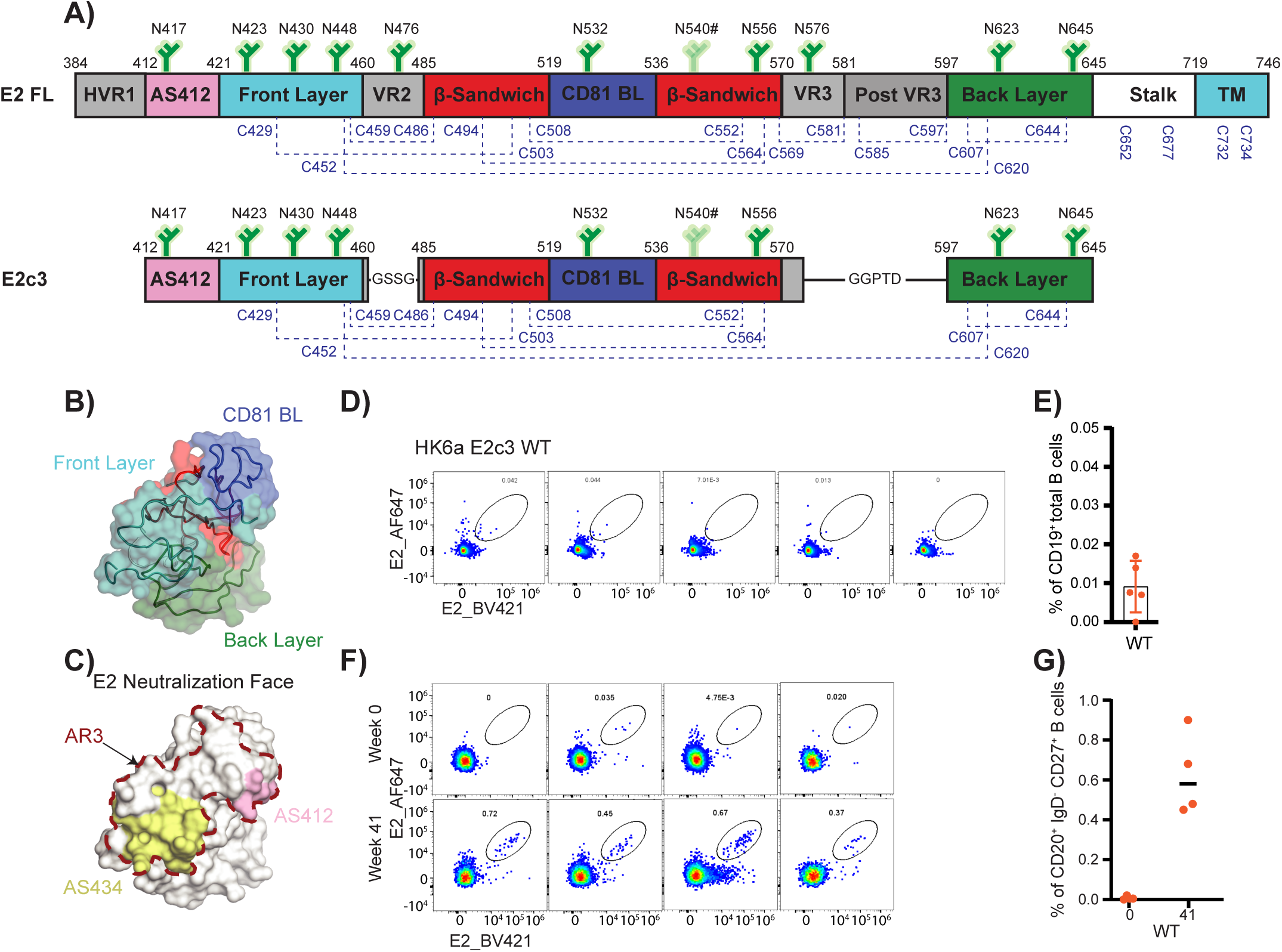
Detection of E2-specific B cells in NBD and NHP immunized with HCV-1 E1E2. **(A)** Schematic representation of the E2 protein with disulfide bridges shown as blue dashed lines and N-Linked glycosylations as green trees. **(B)** The structure of H77 E2c (PDB: 4MWF) with the molecular surface color-coded as in Fig. 1A. **(C)** E2 neutralization face, mapped onto the H77 E2c surface. AR3 is marked in a dashed line, and AS412 and AS434 are colored in pink and yellow, respectively. **(D)** Flow cytometry staining of E2-specific B cells. Cells were gated on CD3^-^ CD19^+^ CD10^-^. **(E)** Frequency of E2-specific B cells in 5 NBDs. **(F)** Flow cytometry staining of E2-specific B cells from NHPs. Cells were gated on CD20+IgD-CD27+. Week 0, pre-immunization; week 41, 1 week after the 4^th^ immunization. **(G)** Frequency of E2-specific B cells in blood samples of 4 NHPs.

Previous failures in clinical vaccination trials (Drane *et al*, 2009; Folgori *et al*, 2006; Cox *et al*, 2019; Page *et al*, 2021) emphasized the need to engineer envelope-derived antigens capable of inducing high-titer bnAb responses. bnAbs isolated from human and animal models (Cowton *et al*, 2018; Keck *et al*, 2018) have provided critical insights into highly conserved epitopes on the HCV E1 and E2 glycoproteins. The vast majority of these bnAbs target E2. Epitope mapping studies reveal that most cross-neutralizing sites, such as antigenic site 412 (AS412), antigenic site 434 (AS434), and antigenic region 3 (AR3), overlap with the CD81 receptor binding site and are highly conserved (Fig. 1C) (Owsianka *et al*, 2006; Kong *et al*, 2013), likely reflecting strong evolutionary constraints on this region, underscoring their relevance for vaccine design. Over the past fifteen years, X-ray crystallography studies of E2-Ab complexes have further defined the antigenic landscape of E2 (Yechezkel *et al*, 2021; Pierce *et al*, 2024; Park *et al*, 2025). These analyses revealed a conserved, conformationally discontinuous, and predominantly hydrophobic surface, termed the E2 neutralization face, that remains accessible to bnAbs despite extensive N-linked glycosylation (Kong *et al*, 2016). This region comprises the front layer (FL) and the CD81-binding loop (CD81 BL) (Fig. 1B, C) and encompasses the principal neutralization sites AR3, AS412, and AS434.

Recently, cryo-electron microscopy structures of the HCV E1E2 heterodimer have provided a structural view of E1 and revealed the overall organization, dynamics, and antigenicity of the native HCV envelope complex (de la Peña *et al*, 2022; Metcalf *et al*, 2023; Augestad *et al*, 2024). These studies show that E1E2 assembles as a dimer of heterodimers (Augestad *et al*, 2024), with interfaces dominated by non-covalent, predominantly hydrophobic interactions. Within this assembly, the E2 stem region is wrapped and stabilized by E1, forming a key structural scaffold. The E2 stem (residues 646–704) packs against the back layer of the E2 core and interfaces extensively with E1. It also contains a highly conserved, E1E2-specific neutralization epitope targeted by the bnAb AR4A (Law *et al*, 2007). Importantly, these structures further demonstrate that the E2 neutralization face remains exposed within the native E1E2 complex, supporting its accessibility to broadly neutralizing Abs, justifying its use as a vaccine antigen.

AR3 consists of a cluster of discontinuous epitopes and is targeted by numerous highly potent cross-genotype bnAbs. Genetic analysis indicates that AR3-specific bnAbs are predominantly, though not exclusively (Ogega *et al*, 2024; Nguyen *et al*, 2025), encoded by the IGHV1-69 (V_H_1-69) heavy-chain variable germline (GL) gene family (Law *et al*, 2007; Giang *et al*, 2012; Merat *et al*, 2019, 2016; Bailey *et al*, 2017; Keck *et al*, 2011, 2019). Shared features across all these V_H_1-69 derived Abs include an atypically hydrophobic heavy chain (HC) complementarity-determining region 2 (CDRH2) (Chen *et al*, 2019), an elongated CDRH3 loop of 15-22 a.a. (compared with 9-15 a.a. for healthy donors) (Briney *et al*, 2019; Tzarum *et al*, 2019), HC-focused interactions (Kong *et al*, 2013), and relatively low levels of somatic hypermutation (SHM) (Kong *et al*, 2013; Tzarum *et al*, 2019, 2020; Flyak *et al*, 2020, 2018). Although they share these features, the E2-bnAb complexes demonstrate different modes of binding, which allows V_H_1-69 encoded bnAbs to be clustered into sub-groups based on their angle of approach to E2 and differences in the footprint on E2 (Yechezkel *et al*, 2021; Chen *et al*, 2020). The difference in the angle of approach can also have a knock-on effect on HC dominance (Chen *et al*, 2019; Tzarum *et al*, 2020). Furthermore, research on B-cell responses in rhesus macaques (RM) vaccinated with the Chiron E1E2 vaccine indicated that AR3 can be targeted by bnAbs encoded by V_H_1-69-equivalent GL genes (Chen *et al*, 2020), with structural analyses showing binding features similar to those of human AR3-directed bnAbs (Chen *et al*, 2021; Nguyen *et al*, 2025; Weber *et al*, 2022).

Together, structural and genetic analyses of V_H_1-69-derived bnAbs show that, despite substantial plasticity within AR3, this region can support interactions with bnAbs that vary in CDR sequence and length. This versatility makes AR3 an attractive target for the design of GL-targeted vaccines. Precedence for this strategy derives from HIV-1 vaccine research, specifically targeting the V_H_1-2 GL as the exclusive ancestor of the VRC01 class bnAbs (Jardine *et al*, 2013). These Abs, which typically target the CD4 receptor binding sites, exhibit minimal reactivity to wild-type (wt) HIV-1 envelope glycoproteins in their GL state. To address this limitation, immunogens were rationally engineered to engage naïve B cells, thereby facilitating the maturation of VRC01 bnAbs in humans (Jardine *et al*, 2015; Venkatesan, 2021). Similar GL-targeting approaches have since been applied to other viral pathogens, including Influenza A (Impagliazzo *et al*., 2015; Yassine *et al*., 2015; Boyoglu-Barnum *et al*., 2021), SARS-CoV-2, and Respiratory Syncytial Virus (McLellan *et al*., 2013), underscoring the broader applicability of this strategy.

The strong bias toward V_H_1-69-derived AR3-targeted Abs provides an enticing rationale for GL-targeted vaccine design in HCV, especially given that previous studies demonstrated that V_H_1-69 GL-reverted Abs exhibit considerable binding affinity for E2 (Tzarum *et al*, 2019; Weber *et al*, 2022; Capella-Pujol *et al*, 2023; Flyak *et al*, 2018), yet lower neutralization potency. Here, we biochemically and structurally characterize the binding of a series of GL-reverted Abs targeting AR3 (AR3A, AR3C, U1, and 212.1.1) to the E2 core protein. The crystal structures of the GL reveted Ab-E2 antigen indicated that GL Abs engage E2 in a highly conserved manner that closely mirrors mature Abs recognition, revealing that key features of the neutralizing epitope are already accessible at the germline level. Leveraging these insights, we rationally engineered E2 variants to enhance interactions with GL-reverted Abs. While our optimized designs improved binding affinity, they did not translate into enhanced engagement of naïve B cells, highlighting an important constraint in GL-targeting vaccine design. Together, our results define both opportunities and limitations for eliciting VH1-69–encoded bnAb responses and provide generalizable principles for antigen design.

## Results

### Limited engagement of naïve B cells to HCV E2

To assess the baseline immunogenicity of wt E2, we quantified the frequency of E2-specific B cells within the naïve B cell repertoire of five HCV-negative normal blood donors (NBDs). The binding of primary B cells to the HK6a E2c3 antigen (E2 residues 412–645 with an internal truncation of VR2, VR3, and the removal of N448 and N576 glycosylation sites; Fig. 1A and methods) was evaluated by fluorescence-activated cell sorting (FACS). These analyses revealed minimal binding, with antigen-specific populations consistently representing fewer than 0.02% of total circulating B cells (Fig. 1D, E). We next examined the frequency of E2-reactive B cells in rhesus macaque (RM) samples from animals vaccinated with HCV-1 E1E2 mRNA or protein antigens, which were previously shown to elicit V_H_1-69-equivalent bnAb responses, although at low circulating levels (Chen *et al*, 2020). Consistent with the NBD data, naïve B cells from pre-immune RMs (week 0) exhibited limited binding to HK6a E2c3 (average 0.02%; Figs. 1F and 1 G). In contrast, 1 week after the 4^th^ immunization (week 41), a substantial increase in HK6a E2c3-specific memory B cells (CD20⁺ IgD⁻ CD27⁺) was observed, with frequencies rising to approximately 0.6% (Fig. 1F, G). These data suggest that wt E2c3 can poorly recruit GL-encoded B-cell receptors (BCRs).

### Reversion to GL Abs Compromises the efficacy of AR3-targeted V_H_1-69 Abs

An in-depth functional and structural analysis of how GL Ab precursors recognize their target antigens can provide essential insights into bnAb induction and enable the design of improved antigens (Chen *et al*., 2020). To this end, we chose four AR3-targeting, V_H_1-69-encoded Abs that we previously used to structurally define the E2 neutralization face: the bnAbs AR3A and AR3C, and the nAbs U1 and 212.1.1. These Abs engage AR3 through two different binding modes (Yechezkel *et al*, 2021) (Fig. EV1B), providing a framework for studying how various GL precursors recognize E2. We used the IMGT V-QUEST program (Brochet *et al*, 2008; Giudicelli *et al*, 2011) to infer GL sequences of the HC (V_H_, D_H_, and J_H_) (Fig. EV2A) [due to the HC dominance, the LC was not reverted] (Tzarum *et al*, 2020, 2019). The inferred GL precursors and variants with only the V_H_ gene reverted were synthesized for further analysis (Ab_GL_ and Ab_1-69_, respectively). Note that our previous study indicated low binding affinity of AR3A_GL_ and AR3C_1-69_ to H77 and HK6a E2c3 (Tzarum *et al*, 2019), therefore, these Abs were excluded from this study. Sequence alignment showed high overall homology between the inferred GL and mature Abs, with 78.12–88.8% amino acid identity (Fig. EV2B). Except for 212.1.1, the most notable differences were localized to CDRH1 (33-50% identity), while CDRH2 showed higher conservation (65–76% identity), including conservation of key hydrophobic residues thought to be involved in interactions with the hydrophobic E2 neutralization face (Fig. 2A).

**Fig. 2:**
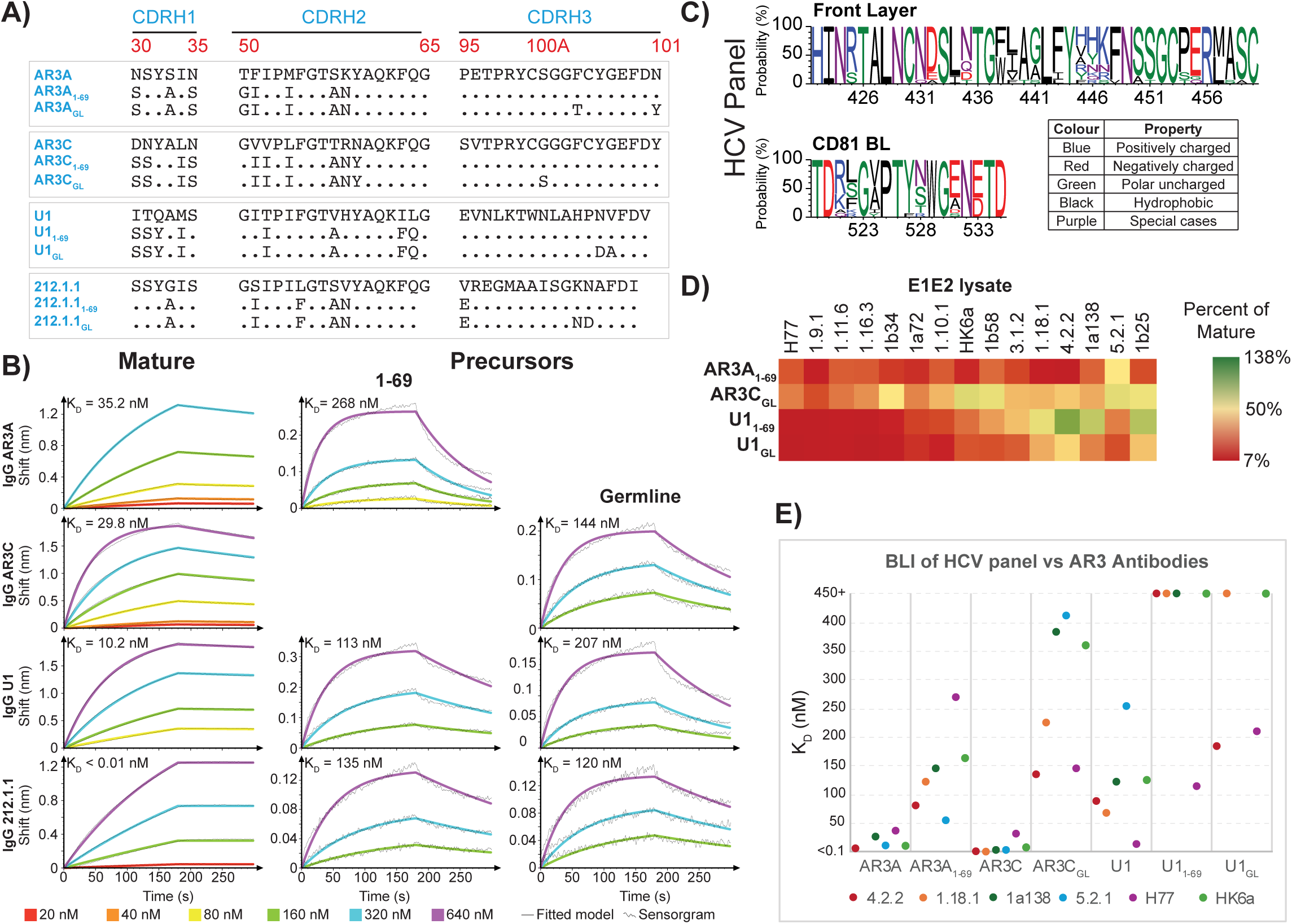
Biochemical characterization of the GL-reverted AR3-targeting Abs. **(A)** Sequence alignment of the HC CDRs of the mature and inferred GL Abs (inferred GL and reversion of only the VH gene to the V_H_1-69 gene sequence) of the AR3-targeting nAbs AR3A, AR3C, U1, and 212.1.1. **(B)** Biolayer Interferometry (BLI) binding kinetics of H77 E2c3 to the mature and GL-reverted Abs. Sensorgrams were obtained from six two-fold titration series starting at 640 nM, and K_D_ values were calculated using a 1:1 global fitting model (solid line). **(C)** LOGO graph presenting the conservation of the front layers and the CD81 BL region of the HCV reference panel. **(D)** ELISA-based cross-reactivity analysis of mature and GL-reverted Abs against E1E2 cell lysate from a reference panel of HCV isolates. Data are presented as a heat map, where binding of GL-reverted Abs is expressed as a percentage of the corresponding mature Ab for each isolate. The color scale is shown on the right. (**E)** Binding affinity (K_D_) summary of mature and GL-reverted Abs against purified E2c3 of UKNP 4.2.1, UKNP 1.18.1, UKNP 5.2.1, 1a138, HK6a, and H77 isolates determined by BLI.

To measure antigen recognition by GL-reverted Abs, we expressed both mature and GL-reverted Abs as IgGs and used biolayer interferometry (BLI) to determine binding kinetics to H77 E2c3 (Fig. 2B). Mature Abs bound E2 with high affinity, with dissociation constants (K_D_) ranging from <0.01 nM for 212.1.1 to 35.2 nM for AR3A. Conversely, GL-reverted variants showed significantly lower affinities, with K_D_ values from 113 nM (U1_GL_) to 268 nM (AR3A_1-69_). The degree of affinity reduction varied greatly among the Abs, from a modest 5–7-fold decrease for AR3A and AR3C to over 1,000-fold for 212.1.1. This reduced binding was mainly due to faster dissociation rates (k_off_), emphasizing the importance of somatic hypermutation in stabilizing the E2-Ab interaction and indicating that GLs have limited ability to maintain stable engagement with E2.

To evaluate the impact of the SHMs on binding breadth and affinity, we compared the binding profiles of mature and reverted GL Abs against native E1E2s from an antigenically diverse HCV reference panel (Salas *et al*, 2022) using enzyme-linked immunosorbent assays (ELISAs). Sequence alignment of the front layer and the CD81 BL regions of the reference panel shows overall high conservation (Fig. 2C). Binding was assessed for mature IgGs (AR3A, AR3C, U1) alongside their respective precursors (AR3A_1-69_, AR3C_GL_, U1_1-69_, and U1_GL_) (Fig. 2D). Initial screening against the 15-panel isolates revealed that, while for some isolates, the precursor Abs showed binding levels more comparable to their mature Ab counterparts, the overall binding magnitude of both mature and GL Abs was lower than that of H77 and HK6a. Consistent with the negligible binding observed in our primary B cell FACS assays, HK6a E1E2 (Genotype 6a) showed a significant loss of recognition upon GL reversion, strengthening the importance of the SHMs in maintaining affinity.

We identified a subset of isolates, including 1b25, UKNP 5.2.1, 1a138, UKNP 4.2.2, and UKNP 1.18.1, that showed the most consistent binding between mature and GL Abs. This suggests that, although the AR3 Abs generally rely on somatic maturation for high-affinity binding, some HCV isolates naturally tolerate GL-encoded Abs more readily, thereby increasing their potential for future vaccine development. To investigate this further, we expressed and purified E2c3 constructs for these isolates (noting that 1b25 E2c3 failed to express) (Fig. EV2C). Subsequent ELISA analysis with these purified antigens confirmed that the isolates maintained comparable binding to the mature and GL Abs, though, as before, the absolute binding was considerably lower than that observed for H77 and HK6a (Fig. EV2D).

To quantify these interactions, we measured the binding kinetics of the purified E2c3 subset, HK6a, and H77 against mature and GL Abs using BLI (Figs. 2E, EV2E, and EV2F). Both AR3A and AR3C exhibited high affinities, with K_D_ values ranging from <0.01 nM (AR3C against UKNP 1.18.1) to 30-35 nM (against H77). The reverted counterpart, AR3A_1-69_, showed consistently higher affinity for the subset isolates (K_D_ values ranging from 53 – 146 nM) compared to H77 (268 nM), and HK6a (162 nM), suggesting these isolates could serve as viable initial scaffolds for vaccine design. Notably, UKNP 5.2.1 exhibited an approximately 5-fold increase in affinity for both mature and reverted AR3A Abs compared with H77. However, in contrast to the previous ELISA results (Fig. 2D, EV2D), AR3C_GL_ showed reduced but measurable affinities that were more consistent with the H77 baseline (144 nM) or the HK6a baseline (359 nM), ranging from 136 nM (against UKNP 4.2.2) to 409 nM (against UKNP 5.2.1) (Fig. 2E). U1 also generally demonstrated lower affinities for the subset isolates, with the reverted U1 Abs showing a particularly sharp decline in binding. These results consolidate the preference for using H77 or HK6a as primary design templates, as they offer a more challenging yet representative baseline for inducing somatic maturation.

### High-resolution Crystal Structures of GL Reverted Fab-E2c3 Antigen Complexes

To provide a high-resolution structural basis for the observed GL reactivity, we determined the crystal structures of the HK6a E2c3 core in complex with the Fab domain of four reverted Abs: AR3A_1-69_, AR3C_GL_, U1_1-69_, and U1_GL_ at 2.0, 1.9, 2.7, and 3.3 Å resolution, respectively (see Table S1 for full details). Across all four structures, the electron density maps clearly defined the Ab-antigen interfaces, allowing for a detailed atomic comparison of GL versus mature antigen engagement.

Overall, no substantial redistribution of the angle of approach (Fig. EV3A) or change in the CDRs’ position and conformation was observed (Fig. 3A), and HC dominance was preserved, consistent with the canonical binding mode of V_H_1-69 encoded Abs (Fig. 3B). Buried surface area (BSA) analysis of the E2c3-Fab interfaces, including contributions from the full Fab, individual HC and LC, and the HC CDR loops (CDRH1-3), revealed only modest differences between GL reverted and mature Abs complexes (Fig. 3A, B). Notably, however, AR3C_GL_ showed no measurable contribution to BSA from CDRH2, indicating an apparent loss of CDRH2-E2c3 interactions that are prominent in the corresponding mature complex (Fig. EV3B). Given the well-known role of the hydrophobic CDRH2 motif in V_H_1-69 mediated recognition (Tzarum *et al*, 2020), this observation was unexpected. Therefore, we assume that this difference results from crystal packing constraints rather than from a biologically relevant change in binding, a point further addressed in the discussion.

**Fig. 3:**
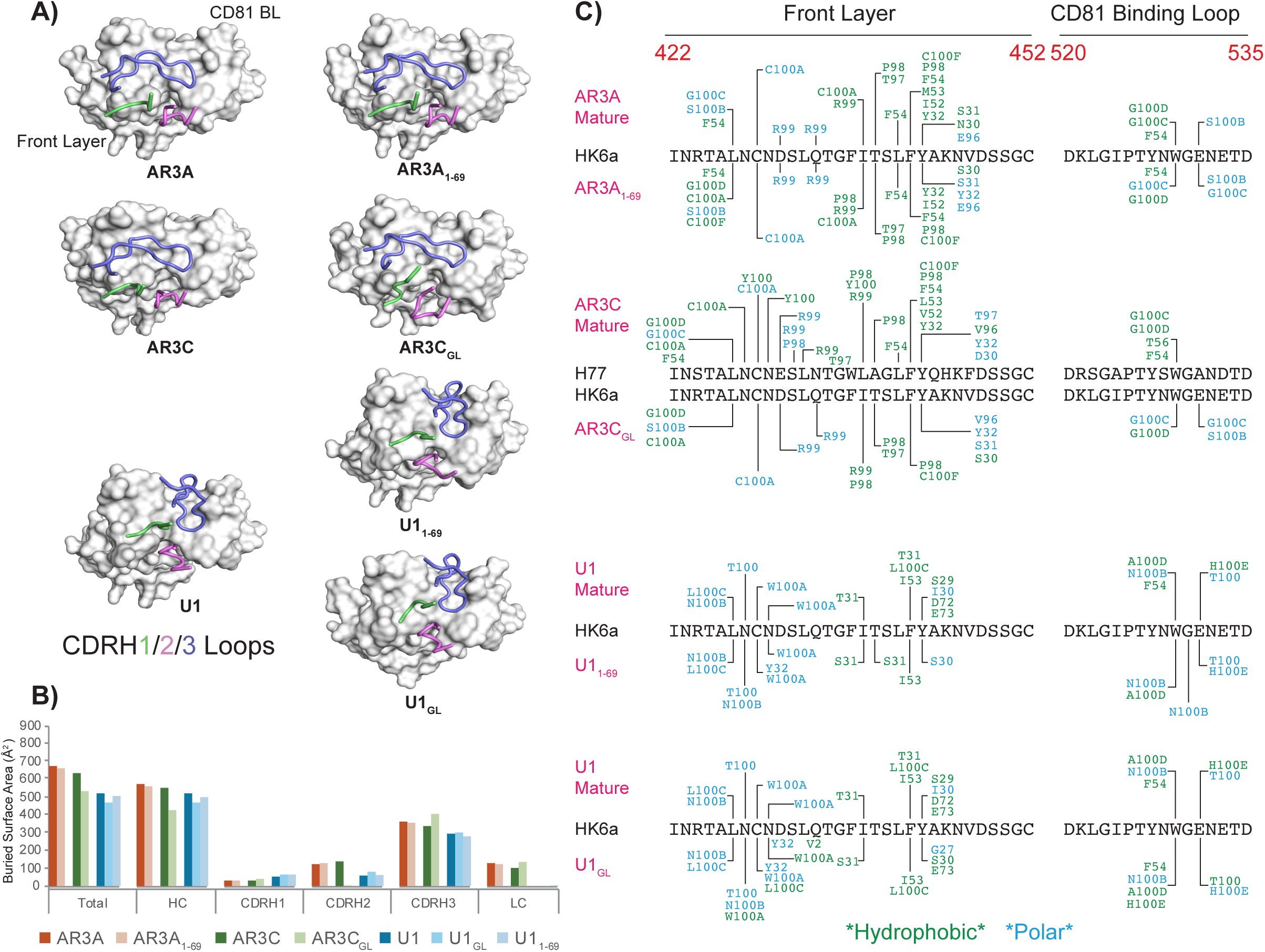
Structural characterization of mature and GL-reverted Abs. **(A)** The position of the CDRH1, CDRH2, and CDRH3 in the E2c3-Fab structures. The E2c3 is shown in surface presentation. The CDRH1, CDRH2, and CDRH3 are shown as cartoons, colored green, violet, and blue, respectively. **(B)** Comparison of BSA between mature complex structures (PDB IDs: 4MWF, 6BKB, 6WO3) and their GL-reverted structures. BSA was calculated for the total Ab-antigen interface, the individual HC and LC, and the specific CDR loops of the HC. **(C)** Schematic contact map summarizing interactions between E2c3 and mature and GL-reverted Abs. The E2c3 front layer and CD81 BL primary sequences are displayed with interactions formed by mature Ab indicated above the sequence and those formed by the precursor Abs indicated below. Hydrophobic interactions are shown in green, and polar interactions are shown in blue.

Next, we mapped the interactions between E2c3 and Abs, focusing on the front layer (a.a. 422-452) and the CD81-binding loop (a.a. 520-535), and compared them with their mature counterparts (Fig. 3C, EV3C). The majority of the interactions were maintained, including contacts mediated by residues that undergo somatic mutation during affinity maturation (e.g. the amino acid at position 30 of the HC, Fig. EV3C). These observations suggest that GL precursors engage the antigen in a manner that closely resembles that of mature Abs.

### Structure-Based Design and Kinetic Validation of Optimized E2c3 Immunogens

Leveraging structural insights from both GL reverted Ab-E2c3 complexes and mature Ab-E2 complexes, we aimed to engineer an E2 antigen with enhanced affinity for GL-reverted Abs. To this end, we employed two design protocols (see Methods) to improve protein stability and enzymatic activity. In the first protocol, we utilized the automated design method FuncLib (Khersonsky *et al*, 2018) to redesign E2 to optimize the Ab-antigen interface, thereby directly strengthening interactions with GL-reverted Abs. FuncLib integrates evolutionary and Rosetta structure-based calculations to identify stable combinations of mutations. The method begins with a phylogenetic analysis to restrict allowable mutations at selected positions to those commonly observed among natural homologs and then models allowed point mutations in Rosetta to eliminate highly destabilizing variants. FuncLib subsequently enumerates and evaluates all permitted mutation combinations using atomistic Rosetta design, followed by relaxation and energy-based ranking.

Given the pronounced conformational flexibility of the E2 front layer (Yechezkel *et al*, 2021; Kong *et al*, 2016), we introduced and successfully expressed five core-stabilization designs (CD1.0-5.0) (Fig. EV4A). However, upon purification, we observed through SEC that these designs lost their tertiary structures. Therefore, we designed two additional iterative designs (CD3.1 and CD3.2) based on CD3, given the shared mutations across all designs. CD3.1 and CD3.2 were expressed and eluted from an SEC column at the same volume as E2c3 wt. Subsequently, CD3.2 was used for downstream BLI and FACS experiments (Fig. 4A, EV4B, EV4C). We also designed five interface-optimized variants (ID1.0-5.0) along with six iterative designs (Fig. EV4A, B). We successfully expressed seven of these designs, but only five retained native folding as confirmed by SEC. Following affinity screening (Data not shown), ID3.0 was identified as the top candidate from this group (Fig. 4A, EV4B).

**Fig. 4:**
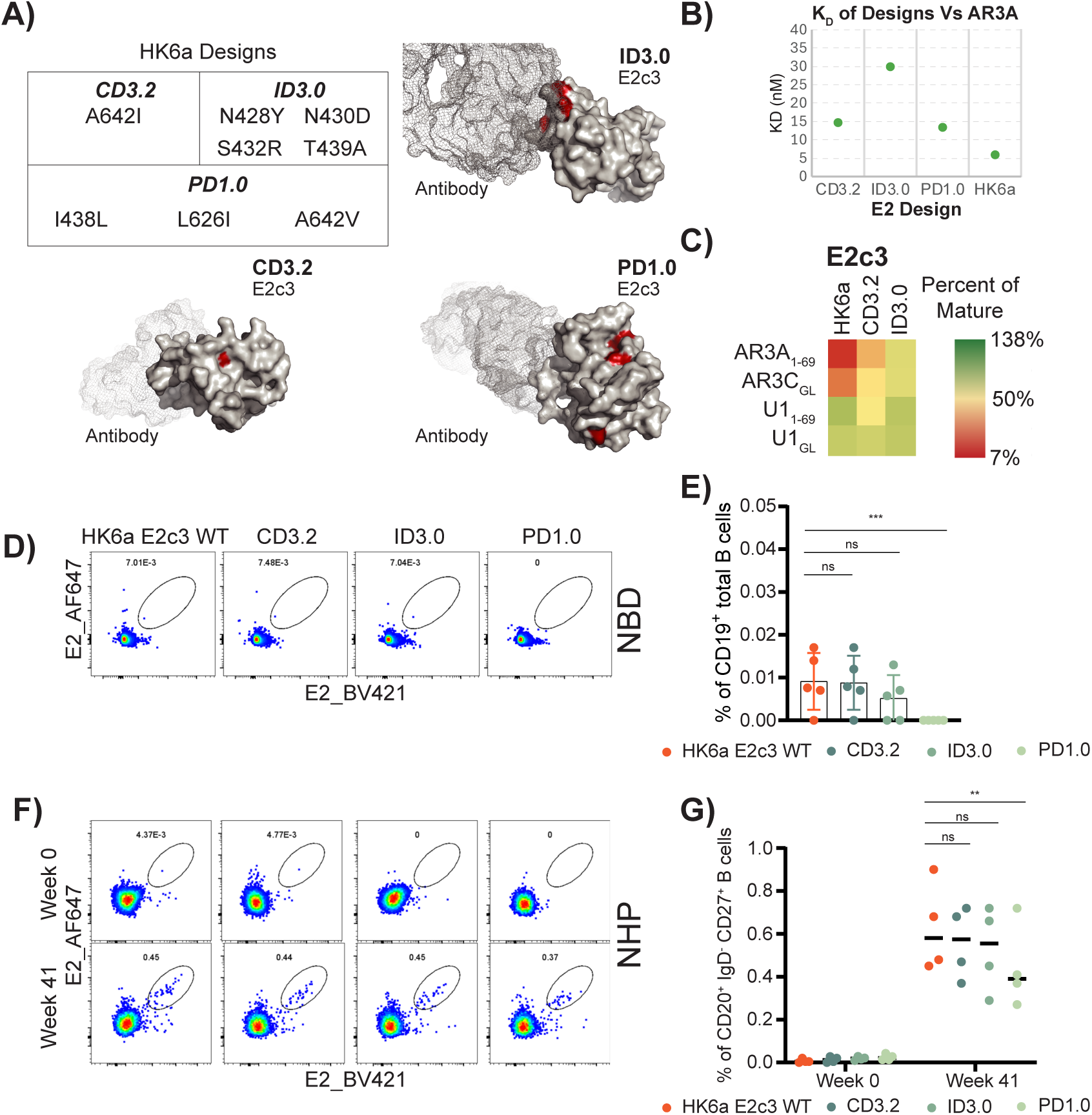
Biochemical Characterization of E2 Design Candidates. **(A)** Structural models of E2 design based on core mutations CD3.2, interface mutations ID3.0, and PROSS-guided stability mutations PD1.0. Mutation loci are highlighted in red on the surface representation of the protein. **(B)** BLI kinetics of the E2 designs and HK6a with IgG AR3A. **(C)** ELISA-based cross-reactivity heat map of the HK6a E2c3 wt, CD3.2, and ID3.0 designs against mature and precursor Abs. Values represent precursor Ab binding as a percentage of the corresponding mature Ab binding, with the color intensity scale indicated on the right. **(D-G)** Detection of E2-specific B cells in NBD and non-human primates (NHP) immunized with HCV-1 E1E2. Week 0, pre-immunization; week 41, 1 week after the 4^th^ immunization. **(D, F)** Flow cytometry staining of HK6a E2c3 wt, CD3.2, ID3.0, and PD1.0 designs specific B cells. Cells were gated on CD20+IgD-CD27+. **(E, G)** Frequency of HK6a E2c3 wt, CD3.2, ID3.0, and PD1.0 designs specific B cells in 5 NBDs and 4 NHPs. Statistical significance was determined using two-tailed Mann-Whitney U-tests. **P < 0.005; ns, not significant.

In the second protocol, we employed the Protein Repair One-Stop Shop (PROSS) method (Goldenzweig *et al*, 2016), which integrates structural and evolutionary information derived from multiple-sequence alignments to introduce mutations that enhance protein stability. PROSS has been widely used to improve enzyme stability and to increase the expression of functional proteins. Notably, PROSS has also been applied to the design of antigens to enhance protein stability, including the HIV-1 envelope gp140 and the *P falciparum* invasion protein RH5 (Campeotto *et al*, 2017; Malladi *et al*, 2020), and the designed variant recently entered clinical trials. More recently, Williams *et al*. used PROSS to engineer stabilizing mutations in the SARS-CoV-2 Spike protein, resulting in improved protein expression and stability while preserving the exposure of neutralizing epitopes. Vaccination studies in mice demonstrated enhanced cross-neutralizing antibody titers elicited by the PROSS-engineered Spike antigen (Williams *et al*, 2023).

Therefore, we generated three cumulative PROSS designs incorporating nested mutations to increase core rigidity and stabilize the overall antigenic complex (PD1.0-1.2) (Fig. EV4A) and successfully expressed and purified them (Fig. EV4B). Of these designs, PD1.0 (Fig. 4A, EV4A) yielded the highest and most homogeneous protein yield and therefore was chosen for downstream functional quantification.

To quantitatively assess the impact of these mutations on AR3A recognition, we performed BLI experiments (Fig. 4B, EV4C). The interface design, ID3.0, exhibited a binding affinity (K_D_) of 29.5 nM, representing a decrease in affinity compared to the 5.1 nM observed for the wt HK6a E2c3. In contrast, the core-stabilization design, CD3.2, and the PROSS-optimized candidate, PD1, yielded K_D_ values of 14.4 nM and 13.3 nM, respectively, affinities closer to the wt HK6a baseline. Subsequent ELISA experiments revealed improved binding of the CD3.2 and ID3.0 designs to the AR3A, AR3C, and U1 germlines compared with their mature counterparts, relative to the HK6a baseline (Fig. 4C). Based on these favorable biochemical and kinetic profiles, these designs were selected as the primary candidates for downstream antigen-specific B-cell sorting and FACS-based assessment of naïve B-cell recruitment.

### Functional Assessment of B-Cell Recruitment by Optimized E2c3 Variants

To determine whether our structure-based designs could enhance immune recognition of the HCV E2 antigen, we evaluated their ability to recruit antigen-specific B cells *ex vivo*. We repeated the B cell binding experiments in both NBDs and RMs immunized with HCV-1 E1E2 using the optimized designs, CD3.2, ID3.0, and PD1, and compared their reactivity against the HK6a E2c3 wt (Fig. 4D-G). We observed no significant differences in naïve B-cell engagement between the wt HK6a E2c3 and the engineered variants in either NBDs or pre-immunized RMs (week 0), with all probes exhibiting negligible baseline reactivity. However, following immunization, the designs CD3.2 and ID3.0 showed E2-specific B cell frequencies comparable to those of the wt antigen in RMs at week 41, one week after the 4^th^ immunization of HCV-1 E1E2. Surprisingly, the PROSS-optimized design PD1.0 exhibited a significant decrease in B-cell binding compared to the wt (P < 0.01) (Fig. 4G). These findings indicate that while our computational designs improved the biochemical binding affinity to mature Abs *in vitro*, these improvements did not translate to enhanced B-cell recruitment in the NHP model. This discrepancy suggests that biochemical affinity alone may be insufficient to overcome the deep-seated GL-binding barriers inherent in the V_H_1-69 lineage Abs.

## Discussion

The clinical observation that approximately 30% of individuals spontaneously clear acute HCV infection provides strong evidence that a protective vaccine is biologically attainable (Hepatitis C, WHO). However, the high rate of viral persistence and the failure of natural immunity to reliably prevent reinfection, particularly among high-risk groups such as PWID, emphasize the urgent need for a vaccine capable of eliciting potent, cross-genotype immune responses (Hepatitis C Surveillance, 2022 Hepatitis Surveillance, CDC).

Effective induction of broad and potent neutralizing responses relies on the successful engagement of bnAb-inferred GL BCRs by the priming immunogen (Stamatatos, 2025). Analyses of naïve B cells from healthy donors and rhesus macaques used in vaccination studies showed minimal recognition of the E2 core antigen, indicating poor engagement of naïve germline BCRs (Fig. 1) (Capella-Pujol *et al*., 2023). These results demonstrate that wt E2 is inherently inefficient at recruiting GL BCRs, highlighting a fundamental challenge in HCV vaccine development. GL-targeting strategies aim to overcome this initial activation barrier by designing immunogens with sufficient affinity to recruit rare or low-affinity precursor B cells. This approach has been validated in HIV-1 vaccine research, where rationally designed immunogens, such as eOD-GT8, have effectively primed VRC01-class V_H_1-2 precursors in human clinical trials. These advancements establish a conceptual framework highly applicable to HCV, where a similar obstacle exists between naïve B-cell recruitment and the development of cross-genotype neutralization. Achieving this goal requires more than understanding the structure of mature bnAb-E2 interactions. It demands a detailed understanding of the GL precursors that trigger these responses (Tzarum *et al*, 2019).

AR3 is a key target for GL targeted structure-based design. Its high sequence conservation and direct overlap with the CD81 receptor-binding site make AR3 an ideal focus for guiding the immune response toward epitopes that are less permissive to viral escape. Moreover, the high plasticity of the AR3 enables binding by various Abs through different binding modes (Kong *et al*, 2016; Tzarum *et al*, 2020; Chen *et al*, 2021; Ogega *et al*, 2024; Weber *et al*, 2022). Consistent with this, repertoire analyses of HCV-infected cohorts demonstrate enrichment of AR3-specific V_H_1-69-encoded or V_H_1-69-encoded-like (Weber *et al*, 2022; Chen *et al*, 2021), emphasizing this lineage as a common immunological solution for targeting the E2 AR3 and the neutralization face. Although other GL pathways have been described (Ogega *et al*, 2024), V_H_1-69 derived Abs remain the most frequently observed and structurally validated mediators of broad neutralization, suggesting that the V_H_1-69 scaffold offers a genetically and structurally favorable framework for engaging conserved AR3 epitopes. Accordingly, vaccine strategies that effectively recruit and guide this lineage may increase the likelihood of eliciting broadly protective responses, while preserving the capacity to accommodate additional neutralizing trajectories.

Biochemical and functional studies, conducted by us and others (Fig. 2), showed that AR3-targeted GL-reverted Abs retain measurable, though reduced, binding to E2, with the degree of reduction varying among Abs. They also exhibit significantly decreased neutralization potency and breadth compared to their mature counterparts. Point mutagenesis analyses of mature Abs further revealed that, for some AR3 bnAbs, decreases in binding and neutralization can be caused by just one or two somatic mutations within CDRH1, such as S30 and S31(Chen *et al*, 2021; Weber *et al*, 2022). Simultaneously, binding and neutralization experiments established that the hydrophobic tip of CDRH2 (residues 53 and 54) is essential for E2 engagement and neutralization by AR3-targeted V_H_1-69 encoded bnAbs(Tzarum *et al*, 2020). Overall, these findings identify specific molecular determinants that govern GL-to-mature transitions and provide a mechanistic foundation for rational antigen design. To mechanistically contextualize these functional observations, we examined the structural basis of GL Abs engagement with E2.

Structural analysis of HK6a E2c3 in complex with V_H_1-69 GL reverted Fabs, AR3A_1-69_, U1_1-69_, and U1_GL_ (Fig. 3), revealed a binding mode that aligns with the typical binding pattern of mature V_H_1-69-encoded Abs (the E2c3-AR3C_GL_ complex will be discussed later). These findings align with the low level of somatic hypermutation and the conservation of interacting residues. A detailed comparison of the E2c3-GL and E2c3-mature Fab complexes showed a similar network of interactions at the Ab-antigen interface (Fig. 3C and EV3C), which could be due to the crystal contacts and represents a limitation of X-ray Crystallography. We could not identify a single specific interaction responsible for the decreased binding affinity observed in the GL-reverted Abs. Additionally, although previous mutagenesis studies have shown that reverting CDRH1 residues at positions 30 and 31 to the GL-encoded Serine significantly reduces binding affinity (Chen *et al*, 2021; Weber *et al*., 2022), our structural analyses suggest that the interactions mediated by these residues are largely preserved in the GL-reverted complexes. A plausible explanation for this difference is that sequence variations between the GL and mature Abs affect the conformational dynamics and stability of the CDR loops, thereby influencing binding energetics without drastically changing the static interaction network observed in the crystal structures. These effects may not be captured in the solid-state environment, where crystallographic constraints can mask subtle differences in flexibility, entropy, or transient contacts that influence affinity in solution.

Notably, the structure of the E2c3-AR3C_GL_ complex revealed a loss of interactions between CDRH2 and E2c3 (Fig. EV3B), whereas those mediated by CDRH1 and CDRH3 remained largely unchanged (Fig. 3A, EV3C). V_H_1-69-encoded Abs possess an unusually hydrophobic CDRH2 (Chen *et al*, 2019). Previous biochemical and structural studies have suggested that this hydrophobic motif initiates engagement with the conserved hydrophobic pocket on E2. We therefore propose that the apparent loss of CDRH2 interactions in the AR3C_GL_ complex may result from subtle differences in crystal packing, rather than reflecting a fundamental alteration in the binding mode.

Our study provides the first structural characterization of AR3-targeted GL-reverted Abs in complex with E2. Previously, we determined the structures of GL-reverted precursors of the AS412-targeted Abs AP33 and 19B3 bound to an AS412-derived peptide (Aleman *et al*, 2018). AS412 is a highly conserved linear epitope located between HVR1 and the E2 front layer (Fig. 1A), which includes residues essential for CD81 engagement, such as W420, and is targeted by several well-characterized bnAbs like HCV1, HC33, and AP33 (Yechezkel *et al*, 2021). Although GL-reverted AS412 Abs demonstrated significantly reduced binding affinity, their structures revealed a binding mode and interaction network similar to that of the mature Abs. Overall, these findings further suggest that affinity maturation can enhance binding by altering CDR loop dynamics and stabilizing the paratope.

An essential step in designing GL-targeted vaccines is to engineer an E2 immunogen with increased affinity for GL Abs to better engage naïve B cells. For this purpose, we utilized two Rosetta-based design protocols and applied two complementary methods. The first focused on reengineering E2 to optimize the Ab-antigen interface, thereby directly enhancing interactions with GL-reverted Abs. The second aimed to stabilize the antigen, thereby increasing its rigidity and thus strengthening its interactions with GL-reverted Abs. Our top designs revealed comparable binding ability of the GL Abs relative to that of mature Abs. Though the designed E2 did not engage naïve B cells more effectively, these findings suggest that future iterations should prioritize interface-specific mutations which provided the most substantial influence on affinity. Furthermore, we suggest that additional factors, including glycan shielding, thermostability and the overall rigidity of the Ab-E2 complex, may play critical roles in the E2 engagement of naïve B cells.

Collectively, our findings represent an initial step toward defining the structural determinants governing GL recognition by AR3-targeting Abs. Further structural and biochemical characterization of the interactions between E2 and GL reverted Abs, combined with structure-guided immunogen design and functional validation, will be required to optimize antigen configurations that efficiently engage naïve BCRs. Such efforts will be essential for the rational development of vaccine candidates that can reliably initiate and steer bnAb responses against HCV.

## Methods

### Cloning and Purification of Antibodies

DNA of mature and reverted Ab HC variable regions, together with IgG constant regions or the Fab constant region, were cloned into the pCMVtpa vector for mammalian cell expression. The Ab variable regions were ordered as DNA fragments and cloned into the pCMVtpa vector downstream to the secretion signal via Gibson Assembly.

DNA from Ab clones was transfected into HEK293expi GnTI-cells using 1 µg of DNA per ml of cells at a concentration of 3 x 10^^6^ cells/ml and incubated at 37 °C, 135 rpm, in 8% CO_2_ for 96 hours. Abs were harvested by centrifugation at 500 g, and the media filtered through a 0,22 µm filter. The media was loaded onto a Protein G column using PBS as the binding buffer and eluted with 100 mM Glycine (pH 2.2). Protein-containing fractions were concentrated and injected into a Superdex 200 Increase 10/300 GL column with a buffer containing 100 mM NaCl and 20 mM Tris-HCl (pH 7.5). Ab purification and correct oligomerisation was confirmed by reducing and non-reducing SDS-PAGE and Western blot using HRP-conjugated anti-human Fc Ab. Western blot membranes were developed using ECL.

### E1E2 and E2 Cloning

E1E2 cloning – the E1E2 genes (amino acids 192-745 based on the H77 prototypic strain, UniProt number P27958) of the following isolates were cloned into the pCMV vector for mammalian cell expression: H77 (Aleman *et al*, 2018), HK6a (Gottwein *et al*, 2009), UKNP1.9.1 (Addgene 97374), UKNP1.10.1 (Addgene 97376), UKNP1.11.6 (Addgene 97382), UKNP1.16.3 (Addgene 98188), 1A72 (Addgene 177690), 1A138 (Addgene 177688), UKNP1.18.1 (Addgene 98190), 1B25 (Addgene 177691), 1B34 (Addgene 177692), 1B58 (Addgene 177693), UKNP3.1.2 (Addgene 98215), UKNP4.2.2 (Addgene 98268), and UKNP5.2.1 (Addgene 98385).

E2c3 cloning – the genes of HCV H77 and HK6a E2c3 (E2 residues 412–645 with an internal truncation of VR2, VR3, and the removal of N448 and N576 glycosylation sites) were cloned into the pCMVtpa vector (Tzarum *et al*, 2019). The E2c3 constructs of 1b25, 5.2.1, 1a138, 4.2.2, and 1.18.1, as determined by sequence alignments to the H77 and HK6a E2c3, were ordered as DNA fragments from Twist Biosciences and cloned into the pCMVtpa vector downstream to the secretion signal via Gibson Assembly.

E2c3 AviTag cloning – the AviTag was cloned to all pCMVtpa E2c3 isolate plasmids between the C-terminus of the E2 and the stop codon by Gibson Assembly (Avi tag Seq – GGCCTGAACGACATCTTCGAGGCCCAGAAGATCGAGTGGCACGAG) E2c3 design cloning – Synthetic gene fragments (Twist Bioscience) corresponding to five initial core variants were cloned into the phCMV expression vector using Gibson assembly, replacing the wt E2c3 sequence. Additional clones were created to facilitate downstream purification by adding a C-terminal Strep-tag II to CD1.0–5.0, placed immediately upstream of the stop codon. For the interface-optimized designs, the wt interface was removed from the phCMV-E2c3 construct and replaced with PCR-amplified fragments containing the desired mutations using hybridized primers. Iterative core and interface variants were generated by site-directed mutagenesis.

### E2 Expression and Purification

H77 E2c3 and HK6a E2c3 were expressed in stable HEK293T GnTI^-^ cell lines that constitutively express the E2c3 core domain. Cells were maintained under standard culture conditions, and the protein was harvested from the growing media.

For the expression of E2c3 of other isolates and the designed E2c3, DNA was transfected into HEK 293expi F or HEK 293expi GnTI^-^ cells (depending on downstream experiments) using 1 µg of DNA per ml of cells at a concentration of 3 × 10^^6^ cells/ml and incubated at 37 °C, 135 rpm, in 8% CO_2_ for 72 hours.

E2s were harvested by centrifugation at 500 g, and the media were filtered through a 0.22 µm filter. The media were loaded onto an IgG AR3A-conjugated Protein G column using PBS as the binding buffer and eluted with 100 mM Glycine, (pH 2.2). Protein-containing fractions were concentrated and injected into a Superdex 200 Increase 10/300 GL column with a buffer containing 100 mM NaCl and 20 mM Tris-HCl (pH 7.5). E2 purification was confirmed by SDS-PAGE and Western blot using IgG AP33 as the primary Ab, followed by HRP-conjugated anti-human Fc Ab. Western blot membranes were developed using ECL.

### Strep-tagged E2c3 Variants Purification

Recombinant proteins were expressed in Expi293F cells as described for E2 expression. Harvested culture media were loaded onto a Strep-Tactin affinity column (GE). The column was washed with 10 column volumes of PBS, and the target proteins were eluted using five column volumes of 2.5 mM desthiobiotin in PBS. Protein-containing fractions were concentrated and injected into a Superdex 200 Increase 10/300 GL column in a buffer containing 100 mM NaCl and 20 mM Tris-HCl (pH 7.5).

### E1E2 lysate expression and purification

HCV E1E2 lysates were expressed by transfection in HEK 293T cells. HEK cells fed with DMEM containing 2 % FBS, 1% L-Glutamine were transfected using Polyethylenimine (PEI, Polysciences) at a 3:1, PEI:DNA, ratio (15 µg of DNA for a 15 cm plate, 66% confluency) and incubated for 4 hours at 37 °C with 5 % CO_2_. After incubation, cells were refed with DMEM containing 10 % FBS, 1 % L-Glutamine, and 1 % Penicillin/Streptomycin and incubated for 72 hours. Cells were consequently harvested by Trypsin treatment and pelleted at 400 g for 5 mins at 4 °C. Cells were washed with ice-cold PBS and pelleted again. Washed cells were resuspended in ice-cold lysis buffer containing 150 mM NaCl, 50 mM Tris (pH 7.5), 2 mM EDTA, 0.5% Triton, and protease inhibitors (10-20 μL of lysis buffer per 10^^6^ cells) and incubated for 30 mins on ice. E1E2 lysate was clarified by centrifugation at 20,000 g for 15 mins at 4 °C. Supernatant was aliquoted and stored at –80 °C.

### PBMC Samples

Peripheral blood mononuclear cells (PBMCs) from normal blood donors (NBDs) were isolated from blood samples collected at the Scripps Research Institute (TSRI). PBMCs were processed using Lymphoprep and SepMate tubes (StemCell) according to manufacturer’s instructions.

PBMC samples from rhesus macaques were obtained from prior HCV-1 E1E2 mRNA or protein immunization studies (Chen et al., 2020; Erin et al., unpublished). All animal procedures were conducted in accordance with protocols reviewed and approved by the Institutional Animal Care and Use Committees (IACUC) of Texas Biomedical Research Institute and TSRI. PBMCs were aliquoted and stored in liquid nitrogen.

### E2 antigen probes and flow cytometry analysis

To generate antigen probes for B cell analysis, Avi-tagged E2c3 protein and variants were produced and biotinylated using the BirA biotin–protein ligase kit (Avidity, Aurora, CO) according to the manufacturer’s instructions. Biotinylated antigens were then conjugated to streptavidin–PE (Thermo Fisher Scientific) or streptavidin–BV421 (BioLegend) at a 4:1 molar ratio.

Antigen-specific B cells were analyzed by flow cytometry. Cryopreserved PBMCs were thawed and aliquoted into 96-well U-bottom plates. Cells were first blocked with anti-CD81 antibody (5 μg/mL, BD Biosciences) and Fc receptor blocker (TruStain FcX™, BioLegend), followed by staining with dual-color–labeled antigen probes at 4°C for 30 min. Cells were then incubated with a cocktail of surface markers for an additional 30 min at 4°C in the dark. Dead cells were excluded using 7-AAD, and samples were acquired on a Cytek Aurora equipped with five lasers using SpectroFlo software (Cytek Biosciences). Data were analyzed using FlowJo (BD Biosciences, version 10.9.0). Gating strategies are shown in Fig. EV1A.

### Biolayer Interferometry (BLI)

Binding affinities were measured using biolayer interferometry (BLI; Gator™). Assay conditions were first optimized to determine the appropriate loading of purified IgGs on anti–protein G biosensors, achieving ∼70% sensor binding capacity. BLI measurements were performed in 96-well plates with the following buffer and sample layout: Column 1 (C1), PBS (baseline); C2, IgG loading; C3–C5, PBS; C6–C12, antigen at 0, 20, 40, 80, 160, 320, and 640 nM.

BLI assays were conducted using protein G probes with the following protocol: 60 s baseline (C1), 60 s IgG loading (C2), 60 s baseline (C3), 180 s association (C6–C12), and 120 s dissociation in PBS (C4 for 0 nM antigen; C5 for antigen-containing samples). The equilibrium dissociation constant (K_D_) was calculated as k_Off_/k_On_ All measurements were performed at 25 °C. Binding data were reference-subtracted against the 0 nM antigen control, and local fitting and kinetic analyses were performed using the Gator software.

### Enzyme-Linked Immunosorbent Assay (ELISA)

Half-well high-binding plates were pre-coated with Galanthus nivalis lectin (GNL, Sigma-Aldrich) at 2.5 µg/mL in PBS to enable enrichment of glycosylated E1E2 from cell lysates. Plates were blocked with 5 % Bovine Serum Albumin (BSA) in PBS. After blocking, plates were washed with PBS containing 0.05% Tween-20 (PBS-T) and incubated with a two-fold serial dilution of E1E2 lysate (from 1:1 to 1:256). Next, plates were washed with PBS-T and tested with IgG AR3A diluted in PBS-T containing 1 % BSA to determine the optimal dilution for 70 % binding. This dilution was used for subsequent experiments with the reverted Abs. Anti-Human Fc conjugated to HRP was used as a secondary Ab diluted in PBS-T 1:20000. Detection was achieved using ECL and quenched with 2 M Sulfuric acid after 20 minutes. Absorbance was measured at 450 nm. For pure E2c3, E2c3 was bound directly to the plate (no GNL) by incubation for one hour at 25 °C. All incubation steps were performed at 25 °C for 1 hour or at 4 °C overnight, except for lysate binding, which was performed at 4 °C overnight. All primary Abs were added at a concentration of 5 µg/ml. Reverted Abs ELISAs were performed on the same plate as their mature counterparts and were normalized against the mature Abs. Significant signal for the mature Abs was defined as a minimum five-fold increase in absorbance relative to the negative control. Binding of the reverted Abs was then expressed as a percentage of the corresponding mature Abs signal, allowing for direct comparison across lineages.

### X-ray crystallization

Crystallization trays were set up using the sitting drop vapor diffusion method at 20°C. Crystallization screen plates were set up using the Mosquito crystallization system (SPTLabtech) with a protein concentration of 10 mg/ml and JCSG I-IV crystal screens as the reservoir conditions. Crystals of HK6a E2c3-AR3A_1-69_ complex were obtained using a reservoir solution of 0.2 M sodium sulfate and 20% (w/v) PEG 3350. Crystals of HK6a E2c3-AR3C_GL_ complex were obtained using a reservoir solution of 0.1 M HEPES (pH 7.5), 20% (w/v) PEG 4000, and 10% (v/v) isopropanol. Crystals of HK6a E2c3-U1_1-69_ complex were obtained using a reservoir solution of 0.2 M Potassium chloride, and 20% (w/v) PEG 3350. Crystals of HK6a E2c3-U1_GL_ complex were obtained using a reservoir solution of 0.16 M Ammonium sulfate, 0.08 M Sodium acetate (pH 4.6), 20% (w/v) PEG 4000, and 20% (v/v) Glycerol. Prior to data collection, crystals were cryoprotected with 10-15% ethylene glycol or 20% PEG200 (E2c3-AR3C_GL_) and flash-cooled in liquid nitrogen. Diffraction datasets were collected at the European Synchrotron Radiation Facility (ESRF) beamlines ID30-A1 (HK6a E2c3-AR3A_1-69_), ID30-A3 (HK6a E2c3-AR3C_GL_, HK6a E2c3-U1_GL_), and ID23-2 (HK6a E2c3-U1_1-69_), Grenoble, France, at 100 K. The structures were solved by molecular replacement using Phaser, with the E2-mature Ab complex as the search model. Structure refinement and model building were conducted in Phenix (Adams *et al*, 2010), and model building was performed with COOT (Emsley & Cowtan, 2004). Final refinement statistics are summarized in Table S1.

### Buried Surface Area Analysis

Buried surface area (BSA) at the Ab-antigen interface was calculated using solvent-accessible surface area (SASA) measurements performed in PyMOL (The PyMOL Molecular Graphics System, Version 3.1.3.1 Schrödinger, LLC.; Shrake & Rupley, 1973), using a rolling-sphere algorithm with a probe radius of 1.4 Å. The antigen (chain E), heavy chain (H), and light chain (L) were first extracted from the full complex to generate monomeric and pairwise structures. SASA values for the antigen alone were calculated from the isolated antigen structure. To determine chain-specific contributions to antigen binding, SASA values of the antigen were recalculated in the presence of either the heavy chain (E+H complex), the light chain (E+L complex), or the full Ab (E+H+L complex). The BSA contributed by each Ab component was computed as the difference between the antigen SASA in isolation and its SASA in the corresponding complex: BSA = SASA (antigen alone) – SASA (antigen in complex).

Per-residue BSA values for the Ab heavy chain were computed by comparing residue-level SASA values in the free heavy chain and in the antigen-bound complex containing only the heavy chain (E+H). SASA calculations were performed with per-atom surface areas stored in the B-factor field and summed per residue. Residue-level BSA was obtained as the difference between the SASA of each residue in the unbound heavy chain and its SASA in the complex. Complementarity-determining region (CDR) contributions were quantified by summing per-residue BSA values over standard heavy-chain CDR definitions (CDRH1 residues 31–35, CDRH2 residues 50–65, and CDRH3 residues 95–102, including insertion codes).

### Protein Design

Positions in the core of the antigen were determined using the Layer residue selector in Rosetta and designed using FuncLib (Khersonsky *et al*, 2018). Due to the paucity of homologs in certain regions, the calculated Position-Specific Scoring Matrix (PSSMs) in these regions were replaced with a PSSM made using a representative panel of HCV variants (Salas *et al*, 2022). Individual mutations with a PSSM score greater than –2 were modelled on the HK6a E2c3 AR3A_1-69_ and AR3C_GL_ structures, and mutations with Rosetta energy less than 2.5 Rosetta energy units relative to each parental structure were accepted. The resulting sequence space was combinatorially enumerated on both structures, and designs were scored based on the “fuzzy”-logic design approach (Warszawski *et al*, 2014) to bias towards designs that optimized the energies of both structures. Designs were clustered, and five designs with at least six mutations relative to one another were selected for experimental testing.

Positions in the antibody-antigen interface were determined using the InterfaceByVector residue selector in Rosetta. Designs were computed as for the core variants, except that the threshold for allowed mutations was set to 3 Rosetta energy units relative to each parental structure. Designs were clustered, and five designs with at least two mutations relative to one another were selected for experimental testing The PROSS (Goldenzweig *et al*, 2016) web server was used to design mutations based on the HK6a E2c3 AR3A, AR3A_1-69_, and the U1_GL_ structure using default settings.

## Acknowledgements

This work is funded in part by the Israeli Science Foundation (ISF), grants #1600/21 and #1818/21 (to N.T.) and by the United States-Israel Bi-national Science Foundation (BSF), grant #2021-165 (to N.T. and M.L.). We acknowledge the European Synchrotron Radiation Facility (ESRF) for the provision of synchrotron radiation facilities under proposal ID MX2402, MX2502, MX2590, and MX2690 and on beamline(s) ID30-A1, ID30-A3. We thank the staff of the ESRF and EMBL Grenoble for assistance and support in using the jointly operated Structural Biology beamline(s).

I.Y., R.F., S.F., and N.T. designed the project; I.Y., H.M., and J.W. cloned, expressed, and purified the E2c3 antigens, IgGs, and Fabs; F.C. and M.L. conducted FACS experiments; I.Y. and H.T. carried out ELISA binding experiments; I.Y. conducted BLI binding experiments; I.Y. and H.M. conducted crystallization experiments and structure determination; I.Y., A.T., and S.F. conducted E2 design; I.Y., F.C., A.T., S.F., M.L., and N.T. analyzed results and wrote the manuscript with support from all authors. The authors declare no competing interests. All data used to understand and evaluate the conclusions of this research are available in the main text and supplementary materials and the Protein Data Bank (accession codes AAAA, BBBB, CCCC, DDDD). Additional data related to this paper may be requested from the authors.

## Fig. legend

**Supplementary 1:**
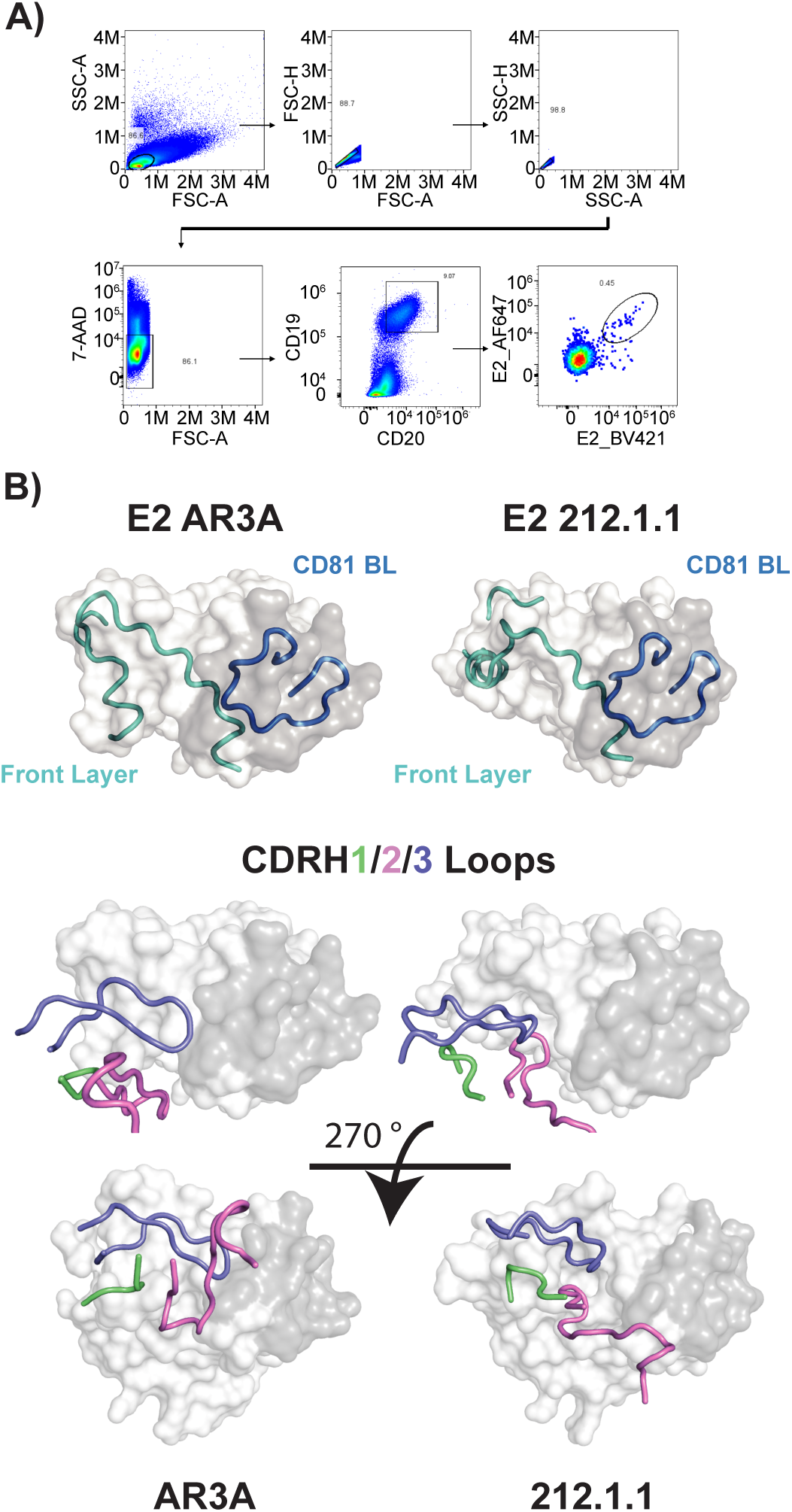
Structural comparison of AR3-series binding modes. **(A)** Flow cytometry gating strategies for analysis of antigen specific B cells. **(B)** Side-by-side view of the E2 complexes for AR3A (PDB ID: 4MWF) and 212.1.1 (PDB ID: 6WO5). Top – the conformation of the front layer. E2 is shown in surface representation with the CD81 BL in grey and the Front Layer in white. The CD81 BL and the Front Layer are further highlighted in cartoon representation, colored in dark blue and cyan, respectively. Bottom – the position of the CDRHs in the E2c3-Fab structures. The CDRH1, CDRH2, and CDRH3 are shown as cartoons, colored green, violet, and blue, respectively.

**Supplementary 2:**
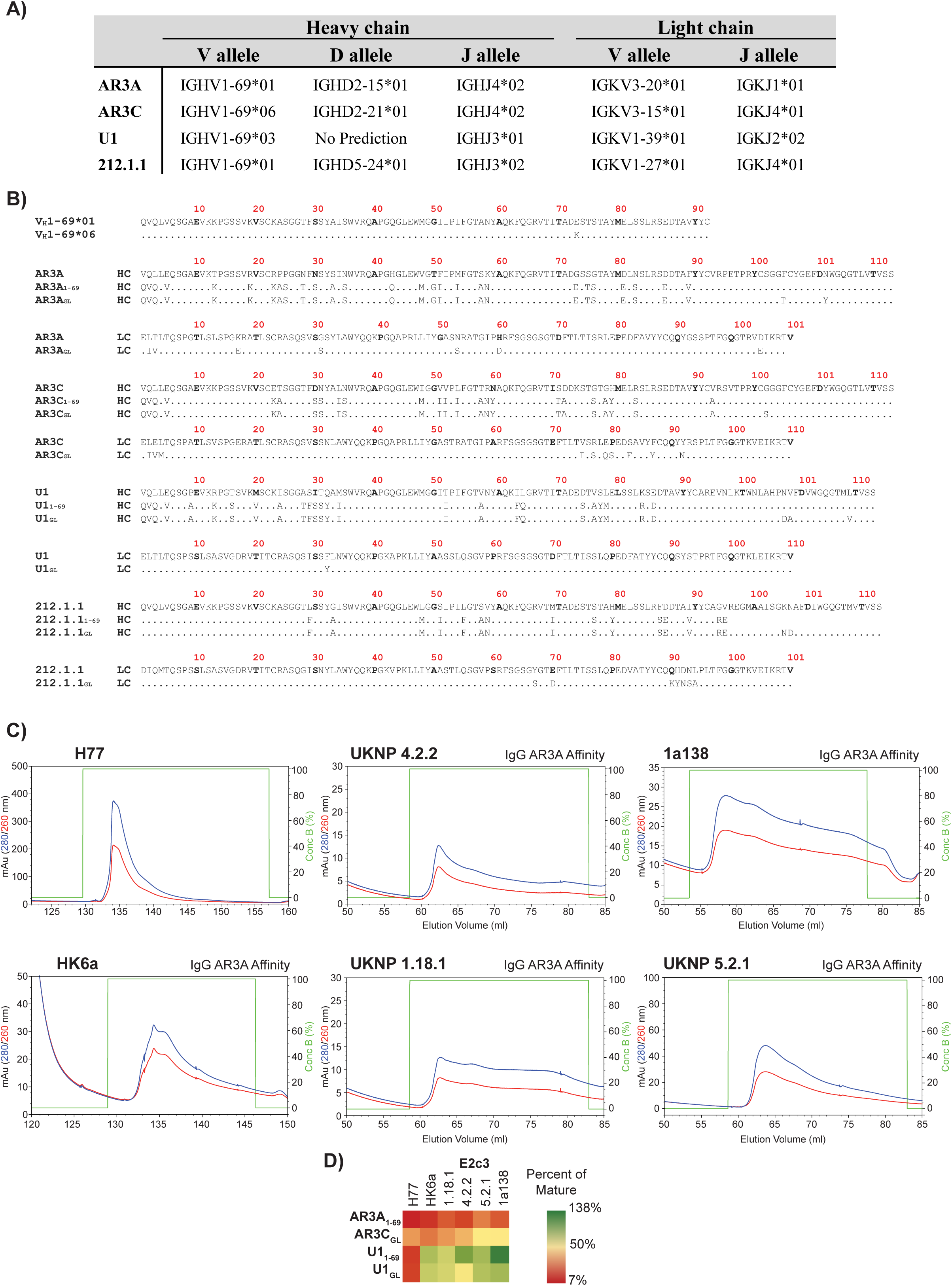
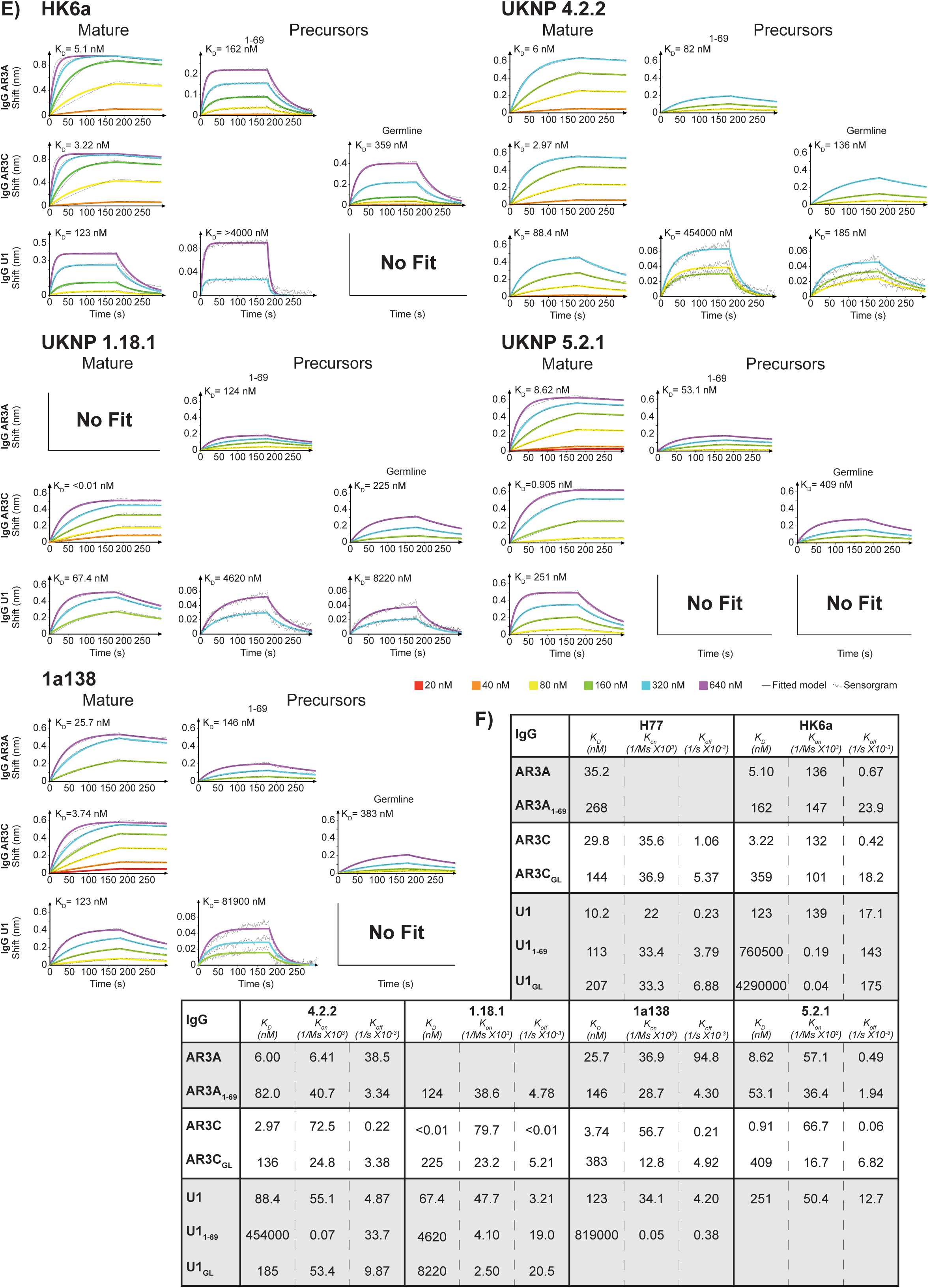
Biochemical characterization of the GL-reverted AR3-targeting Abs. **(A)** IMGT V-QUEST predicted GL alleles of the HC (V_H_, V_D_, and J_H_) for four AR3-targeting, V_H_1-69-encoded bnAbs. Note, due to the HC dominance, the LC was not reverted. **(B)** Multiple sequence alignment of the variable regions of the mature AR3 nAbs and their corresponding GL reversions. **(C)** Affinity chromatography profiles (using IgG AR3A conjugated Protein G beads) for the purification of H77, HK6a, UKNP 4.2.2, UKNP 1.18.1, 1a138, and UKNP 5.2.1. **(D)** ELISA-based cross-reactivity analysis of mature and GL-reverted Abs against E2c3 of H77, HK6a, UKNP 4.2.2, UKNP 1.18.1, 1a138, and UKNP 5.2.1. Data are presented as a heat map, where binding of GL-reverted Abs is expressed as a percentage of the corresponding mature Ab for each isolate. The color scale is shown on the right. **(E)** BLI binding kinetics of HK6a, UKNP 4.2.1, UKNP 1.18.1, UKNP 5.2.1, and 1a138 E2c3 to the mature and GL-reverted Abs. Sensorgrams were obtained from six two-fold titration series starting at 640 nM, and K_D_ values were calculated using a 1:1 global fitting model (solid line). **(F)** Summary table of kinetic constants (K_D_, K_on_, and K_off_) for the interactions of HK6a, UKNP 4.2.2, UKNP 1.18.1, 1a138, and UKNP 5.2.1 E2c3s with mature and GL-reverted Abs, as determined from BLI.

**Supplementary 3:**
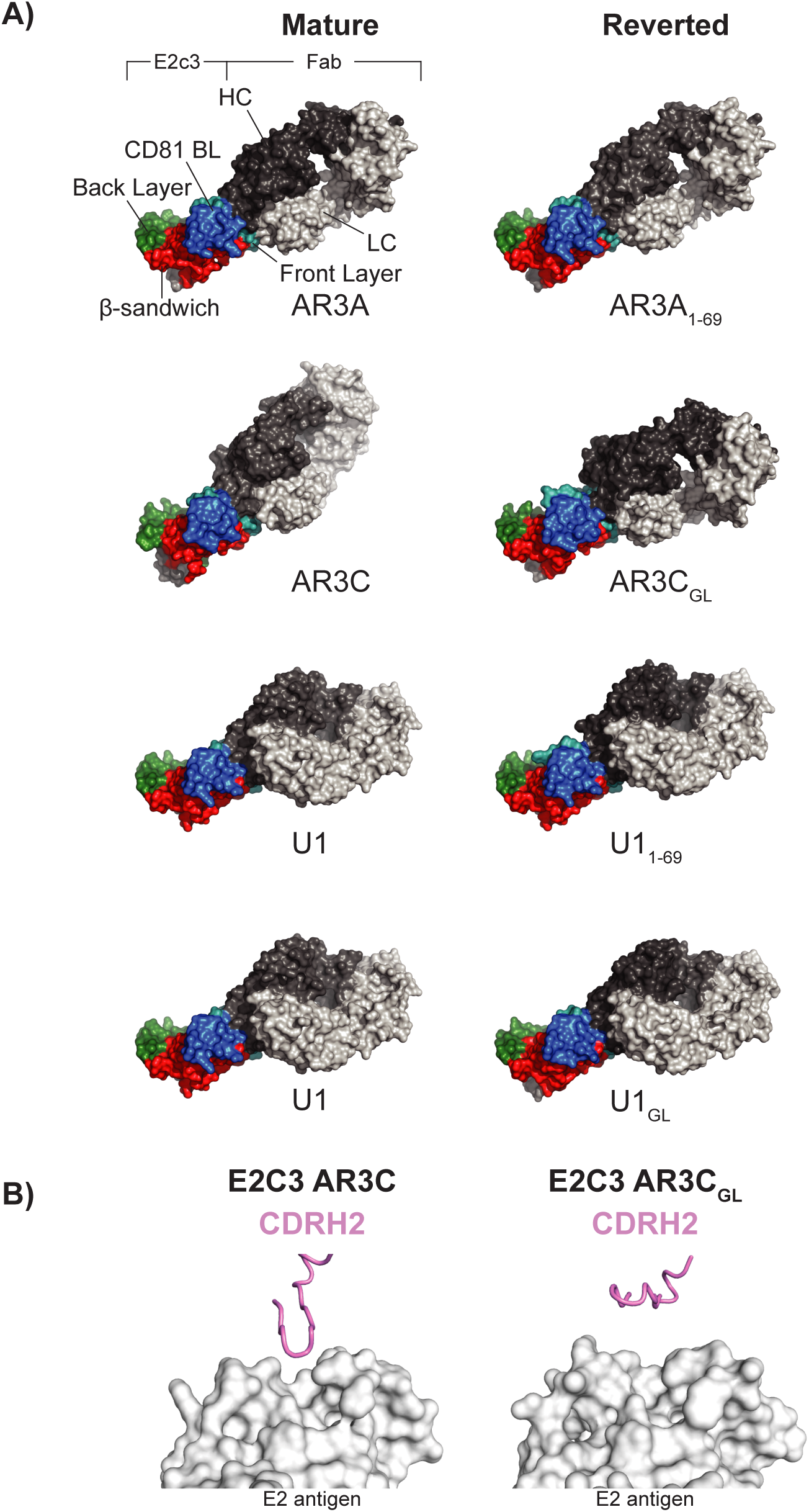
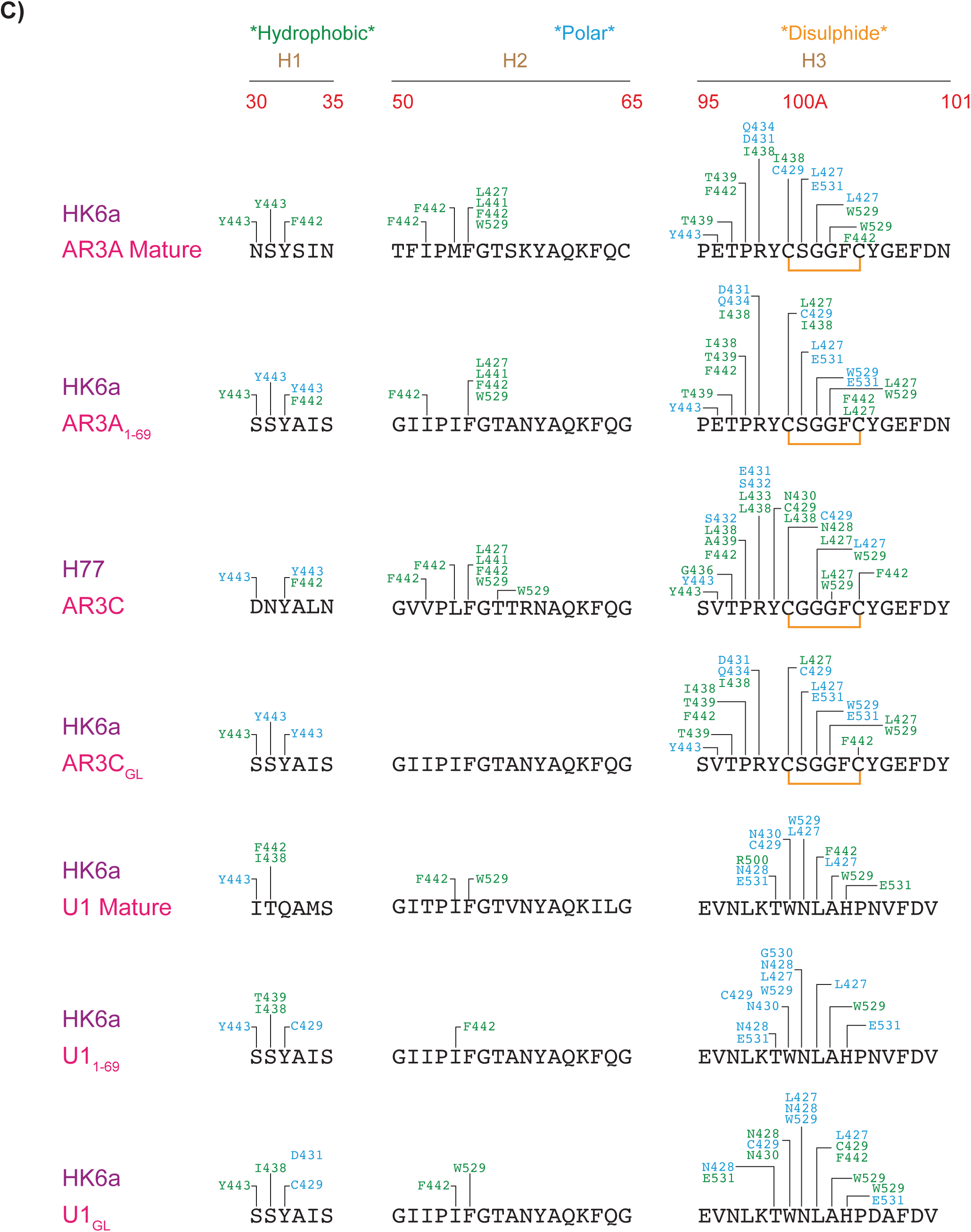
Structural characterization of mature and GL-reverted Abs. **(A)** Surface representation of mature and GL-reverted Ab-E2c3 crystal structures. All structures are superimposed on the E2c3 of the HK6A E2c3-AR3A complex. E2c3 is color-coded by structural domain: front layer in cyan, beta-sandwich in red, CD81 BL in blue, and back layer in green. **(B)** View of mature (PDB ID: 4MWF) and GL-reverted AR3C CDRH2 interactions with E2c3. **(C)** Schematic contact map summarizing interactions between E2c3 and mature and GL-reverted Abs. The CDRH loops primary sequences are displayed with interactions formed by E2c3 indicated above each sequence. Hydrophobic interactions are shown in green, and polar interactions are shown in blue.

**Supplementary 4:**
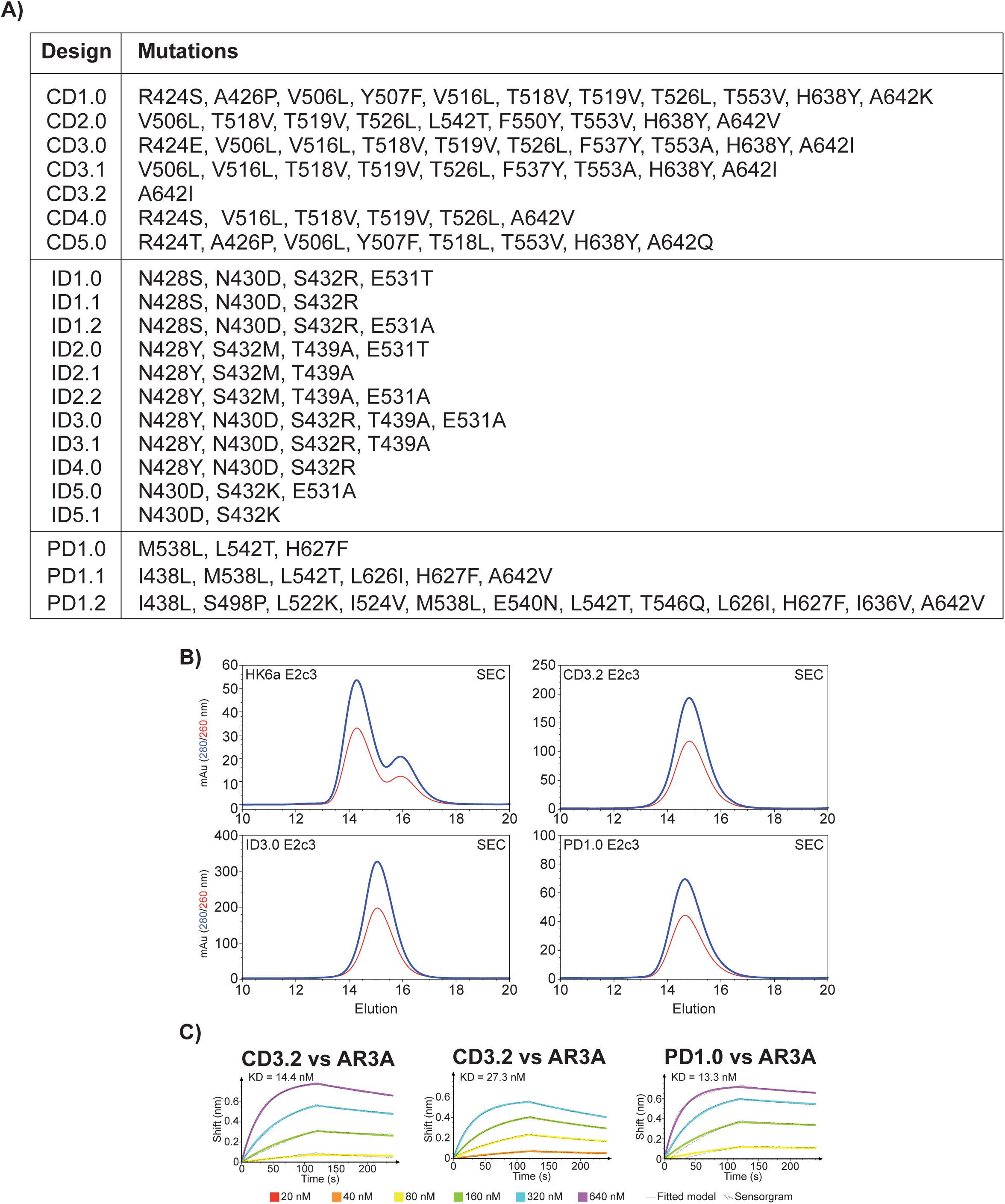
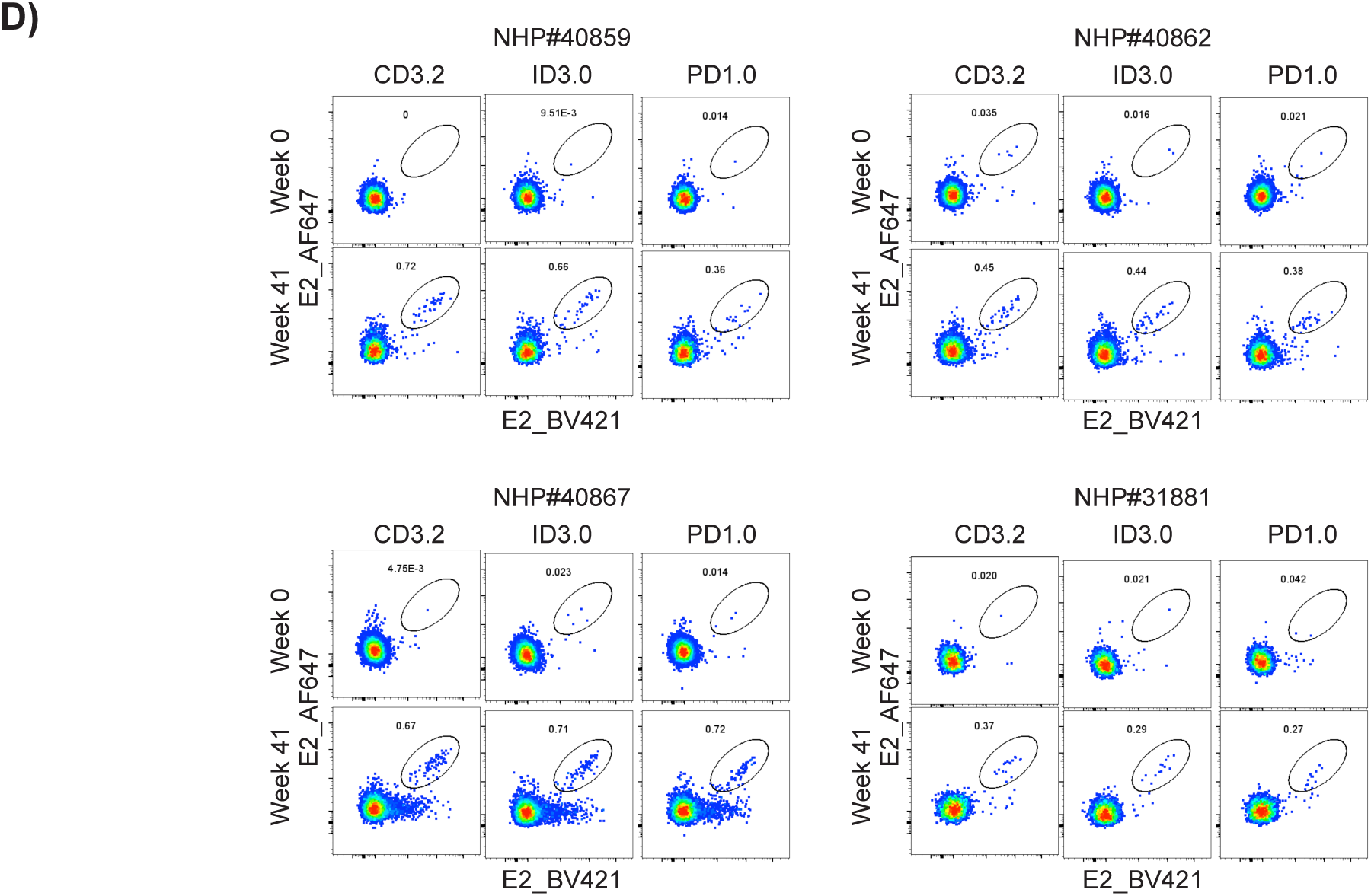
Biochemical Characterization of E2 Design Candidates. **(A)** Table of total E2 design candidates divided by method of design: Core designs (CD), Interface designs (ID), and PROSS designs (PD). **(B)** Size exclusion chromatography profiles of HK6a, CD3.2, ID3.0, and PD1.0 E2c3. (**C**) BLI kinetics of the HK6a E2c3 designs and wt with IgG AR3A.

**Table S1.** Data collection and refinement statistics for HK6a E2c3-AR3A_1-49_, HK6a E2c3-AR3C_GL_, HK6a E2c3-U1_1-49_, and HK6a E2c3-U1_GL_.

| Data collection | HK6a E2c3-AR3A<sub>1-69</sub> | HK6a E2c3-AR3C<sub>GL</sub> | HK6a E2c3-U1<sub>1-69</sub> | HK6a E2c3-U1<sub>GL</sub> |
| --- | --- | --- | --- | --- |
| Beamline | ESRF ID-30A-1 | ESRF ID-30A-3 | ESRF ID-23-2 | ESRF ID-30A-3 |
| Wavelength (Å) | 0.96546 | 0.96770 | 0.8731 | 0.96770 |
| Space Group | P2<sub>1</sub>2<sub>1</sub>2<sub>1</sub> | P2<sub>1</sub>2<sub>1</sub>2<sub>1</sub> | P2<sub>1</sub>2<sub>1</sub>2<sub>1</sub> | P2<sub>1</sub>2<sub>1</sub>2<sub>1</sub> |
| Unit cell parameters (Å, °) | a=61.4, b=66.9, c=218.7 | a=60.3, b=64.4, c=183.3 | a=59.8, b=79.0, c=133.5 | a=60.3, b=82.0, c=135.9 |
| Resolution range (Å) | 46.98-2.04 (2.08-2.11)<sup>a</sup> | 44.05-1.93 (2.00-1.93) | 54.61-3.30 (3.36-3.30) | 45.73-2.75 (2.85-2.75) |
| Observations | 228,946 | 240,241 | 98,274 | 81,061 |
| Unique reflections | 57,518 (2,120) | 54,095 (5,240) | 9,197 (480) | 17,730 (8,068) |
| Completeness (%) | 98.8 (98.7) | 94.8 (95.3) | 92.2 (100) | 95.7 (96.7) |
| $\langle I/\sigma(I) \rangle$ | 7.2 (0.7) | 7.0 (0.8) | 5.1 (0.9) | 8.3 (1.2) |
| CC<sub>1/2</sub><sup>b</sup> | 0.99 (0.47) | 0.99 (0.32) | 0.97 (0.33) | 0.99 (0.46) |
| R<sub>merge</sub><sup>c</sup> | 0.07 (1.09) | 0.12 (1.57) | 0.68 (8.66) | 0.10 (0.93) |
| R<sub>pin</sub><sup>d</sup> | 0.05 (0.93) | 0.07 (0.89) | 0.20 (2.62) | 0.07 (0.69) |
| Redundancy | 4.0 (3.8) | 4.4 (4.6) | 1.7 (1.7) | 4.6 (4.8) |
| Refinement Statistics |  |  |  |  |
| Resolution (Å) | 46.98-2.04 (2.08-2.04) | 44.05-1.93 (2.00-1.93) | 54.61-3.31 (3.42-3.31) | 45.73-2.75 (2.85-2.75) |
| No. reflections | 56,082 | 53,734 | 8,674 | 17,703 |
| R<sub>cryst</sub><sup>e</sup>/R<sub>free</sub><sup>f</sup> | 0.235/0.280 | 0.201/0.245 | 0.268/0.323 | 0.236/0.281 |
| Number of atoms |  |  |  |  |
| Protein | 4,626 | 4,661 | 4,362 | 4,531 |
| Glycan | 70 | 126 | 28 | 41 |
| Water | 206 | 327 | 15 | 22 |
| Average B value (Å<sup>2</sup>) |  |  |  |  |
| All proteins | 66 | 37 | 81 | 72 |
| Glycan | 101 | 65 | 113 | 100 |
| Water | 62 | 46 | 62 | 67 |
| R.m.s. deviation from ideal geometry |  |  |  |  |
| Bond length (Å) | 0.014 | 0.008 | 0.013 | 0.014 |
| Bond angle (°) | 1.2 | 1.2 | 1.6 | 1.6 |
| Ramachandran Plot (%)<sup>g</sup> |  |  |  |  |
| Favored | 96.6 | 97.0 | 95.3 | 95.0 |
| Outliers | 0.3 | 0.6 | 0.9 | 0.5 |
| PDB ID | XXXX | YYYY | ZZZZ | WWWW |

<sup>a</sup> Parentheses refer to outer shell statistics.
<sup>b</sup> CC<sub>1/2</sub> = Pearson Correlation Coefficient between two random half datasets.
<sup>c</sup> $R_{\text{merge}} = \sum_i \sum_h |I_{hi}(i) - \langle I_{hi} \rangle| / \sum_i \sum_h I_{hi}(i)$ , where $I_{hi}(i)$ is the scaled intensity of the $i^{\text{th}}$ measurement of reflection $h, k, l$ , and $\langle I_{hi} \rangle$ is the average intensity for that reflection.
<sup>d</sup> $R_{\text{pin}} = \sum_i (1/(n-1))^{1/2} \sum_h |I_{hi}(i) - \langle I_{hi} \rangle| / \sum_i \sum_h I_{hi}(i)$ , where $n$ is the redundancy.
<sup>e</sup> $R_{\text{cryst}} = \sum_i |F_o - F_c| / \sum_i |F_o| \times 100$ , where $F_o$ and $F_c$ are the observed and calculated structure factors.
<sup>f</sup> $R_{\text{free}}$ was calculated as for $R_{\text{cryst}}$ , but on a test set of 5% of the data excluded from refinement.
<sup>g</sup> Calculated using MolProbity

